# Characterisation of sperm piRNAs and their correlation with semen quality traits in swine

**DOI:** 10.1101/2020.03.16.994178

**Authors:** M. Ablondi, M. Gòdia, J. E. Rodriguez-Gil, A. Sánchez, A. Clop

**Author notes:** These authors contributed equally to this work (M. Ablondi, M. Gòdia).

## Abstract

Piwi-interacting RNAs (piRNAs) are a class of non-coding RNAs which main reported function consists on the silencing of transposable elements and genome stability in mammalian germline. In this study we have identified piRNAs in porcine sperm, using male germline and zygote datasets from human, mice, cow and swine, and evaluated the relation between their abundances and sperm quality traits. Our analysis identified 283,382 piRNAs, 1,355 of which correlated to at least one semen quality trait. Indeed, genome analysis of the correlated piRNAs evidenced that 57% of these were less than 50kb apart and were significantly enriched near Long Interspersed Nuclear Elements (LINEs). Moreover, some of the significant piRNAs mapped within or close to genes relevant for fertility or spermatogenesis such as *CSNK1G2* and *PSMF1*.

## Main text

Fertility is of paramount importance in nowadays’ livestock production since it impacts on the efficiency of the system and thus on its sustainability. Pig production highly depends on the genetic merit of elite boars used for artificial insemination (AI) and on their sperm quality, which is ultimately used to spread the pigs’ genetic material (Zak *et al.* 2017). Several independent studies have shown the involvement of small non-coding RNAs (sncRNAs) on the molecular mechanisms that regulate spermatogenesis, testis development (He *et al.* 2009; Gebert *et al.* 2015; Goh *et al.* 2015) and fertility (Salas-Huetos *et al.* 2015). Piwi-interacting RNAs (piRNAs) are a class of sncRNAs with an average size between 26 and 32 bp of length that are highly abundant in sperm (Gòdia, *et al.* 2018a). piRNAs are crucial for transposon silencing and genome stability during spermatogenesis and early embryo development (Aravin *et al.* 2008; Yan 2017). Studies in mutant Piwi proteins have enlighten their role in spermatogenesis and fertility (Fu and Wang 2014; Gòdia *et al.* 2018a). Few studies have evaluated the relationship between piRNA abundance and semen quality in human (*Cui et al.* 2018) and cattle (*Capra et al*. 2017) but this remains unchecked in swine.

To shed light into the relevance of piRNAs on sperm quality traits in swine, we have apiRNA characterisedthe piRNA composition of the porcine sperm and have evaluated the correlation between piRNA abundance and semen quality traits.

34 fresh sperm ejaculates, each from a different boar, were obtained by specialised professionals and spermatozoa were directly purified from the ejaculate by density gradient centrifugation (Gòdia *et al.* 2018b). Details on RNA extraction and short RNA-seq analysis are described in Gòdia *et al.* (2018b) and Gòdia *et al.* (2019). The fastq files can be found at the SRA (accession number PRJNA520978). Raw reads quality was evaluated with FastQC 0.11.9 (https://www.bioinformatics.babraham.ac.uk/projects/fastqc/).

Low-quality reads, sequence adaptors and reads with length < 26 bp were removed with Cutadapt 2.8 (Martin 2011). Filtered reads were then mapped to the pig genome (Sscrofa11.1) with Bowtie 1 (Langmead 2010), using the “--best--strata” option and allowing for 1 mismatch. Previously annotated piRNAs in relevant tissues (sperm, testis and zygote) in *Sus scrofa, Bos taurus, Homo sapiens* and *Mus musculus* were retrieved from the piRBase database v2.0 (Wang *et al.* 2019). piRNA abundance was quantified using FeatureCounts 1.6.4 (Liao *et al.* 2014) and read counts were normalised by sequencing depth as counts per million (CPM). Only piRNAs with CPM > 1 in at least 75% (N = 26) of the samples were kept. piRNA abundances were then stabilised with a log2 transformation. Pearson correlations between piRNAs and the 13 semen quality traits assessed were calculated (Table S1). Correlations with P ≤ 0.01 were considered significant. Statistical overrepresentation of biological processes of genes present within a ± 2.5kb window from significant piRNAs was conducted with PANTHER 14.0 (http://pantherdb.org) using Human as reference and by setting the level of significance equal to q-value ≤ 0.01.

As average, 2.8 M reads passed the quality control and length filters and were used for piRNA characterisation. Overall, 283,382 piRNAs were successfully mapped to the porcine genome (Table S2) and 6,729 piRNAs had CPM > 1 in more than 75% of the samples. The piRNAs had the characteristic piRNA signature previously described in pig (Gebert *et al.* 2015) and cattle (Rosenkranz *et al.* 2015), of a preference for a uridine (U) at their 5’ end (51.2%) but lacking an enrichment of the adenine (A) at position 10 as found in human (Yang *et al.* 2019) (Figure S1). A total of 1,355 piRNAs were significantly correlated with at least one phenotype at P ≤ 0.01 (Table S3). Significant correlations ranged between −0.72 to 0.64 and the number of associated piRNAs varied considerably across phenotypes and chromosomes (Figure S2, Figure 1a).

**Figure 1.**
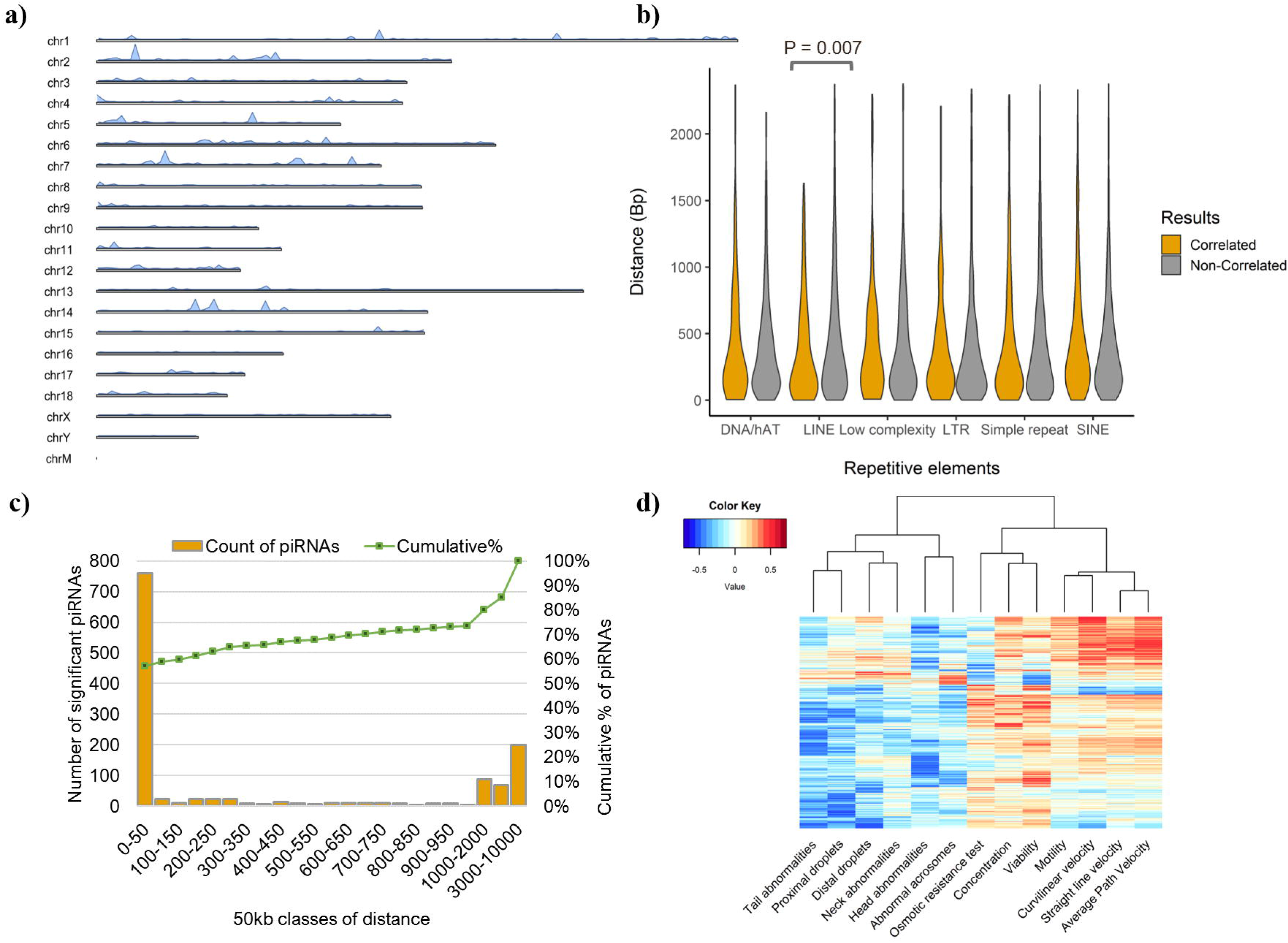
Characteristics of the sperm piRNAs associated to semen quality. **a)** Genome wide location of the piRNAs significantly associated to sperm quality traits. **b)** Violin plot of the distance between piRNAs and Repetitive Elements (RE) for the phenotype-correlated and the non-correlated piRNAs. The significant difference is highlighted with the symbol * **c)** Distance distribution of significant piRNAs across 50kb window size. **d)** Heatmap of the piRNA correlations for the 13 phenotypes. The cells in the plot are coloured according to the Pearson correlation coefficient values with more intense colours indicating higher positive (red) or negative (blue) correlations.

The piRNAs associated to semen quality mapped, on average, less than 500 bp away from a Repetitive Element (RE). Nearly 34%, 25.3% and 21.3% of these piRNAs were close to Short Interspersed Nuclear Elements (SINEs), simple repeats and Long Interspersed Nuclear Elements (LINEs), respectively (Table S4). Interestingly, the piRNAs associated to semen quality traits tended to be closer to LINEs (Wilcoxon rank sum test, *P* = 0.007) compared to the list of piRNAs not correlated with semen quality (Figure 1b). These findings go in line with the potential role of piRNAs in transposon silencing and germline genome stability as reported in several mammalian species (O’Donnell and Boeke 2007) (Figure S3). Di Giacomo et al (2013) found that in particular, piRNAs silence LINE1.and disrupting the silencing of LINEs has been associated to spermatogenesis defects (reviewed by Gòdia *et al.* 2018a).

Previous studies have shown that piRNAs tend to cluster in the genome (Brennecke *et al.* 2007; Capra *et al.* 2017). We evaluated the distance between contiguous piRNAs showing significant phenotypic associations and identified that 57% of these were less than 50kb apart (Figure 1c). Five-hundred twenty-seven of the 1,355 significant piRNAs gathered within 50 clusters with a mean length of 18kb and an average of 10.5 piRNAs per cluster (Liu *et al.* 2012; Li *et al.* 2019).

The piRNA : trait correlations were also analysed with a hierarchical clustering algorithm. In general, semen quality correlations showed that the abundance of individual piRNAs has a beneficial impact on semen quality (Figure 1d). Noteworthy, this general trend was also found when considering the total piRNA abundance (the sum of the CPM values of each individual piRNA) and the semen quality traits (e.g. percentage of tail -TABN- and head abnormalities -HABN-, corr.= −0.32, corr.= −0.31, *P* = 0.07), although none of the correlations reached significance levels (Table S5). Overall, these results indicate that RE silencing and genome stability impacts on spermatogenesis and semen quality as previously suggested (Ernst *et al.* 2017; Ge *et al.* 2017).

A total of 541 protein-coding genes were located within the 5kb region centred at each piRNA displaying significant correlation with semen traits. Gene ontology analysis showed enrichment for 11 biological processes, including reproduction or positive regulation of cell projection organisation (Table S6). Some of these genes, *CATSPER2, CATSPERG, OAZ3, ODF1, ODF2, PRM1, TEX14, TSSK2, TSSK3* and *TSSK6* are known to play key roles in spermatogenesis, sperm hyperactivation, sperm morphology, and male fertility (Tokuhiro *et al.* 2009; Waclawska and Kurpisz 2012; Yang *et al.* 2012; Zhang *et al.* 2012; Savadi-Shiraz *et al.* 2015; Cai *et al.* 2017; Sun *et al.* 2017).

Eight of the 10 piRNAs most significantly correlated with semen quality traits, presented negative correlation values (Table 1a). Once more, this suggests that a reduction of piRNA abundance leads to a decrease of semen quality. This could be due to a deficient silencing of REs and the consequent increase of genome instability which could impact germ cell correct development (Di Giacomo *et al.* 2013; Gunes and Kulac 2014). Some of these 10 piRNAs located near genes with functions related to spermatogenesis. A positive correlation was found between piR-ssc-112967 and the Osmotic Resistance Test quality trait (ORT) (corr. = 0.62). This piRNA maps within the *CSNK1G2* gene. *CSNK1G2* has been linked to sperm surface modifications, sperm maturation and sperm–egg communication in bull sperm (Byrne et al. 2012). piR-ssc-113649, which correlated with the percentage of viable cells (VIAB), abnormal acrosomes (ACRO) and TABN (corr. = 0.64, −0.51 and −0.50, respectively), maps within the *PSMF1* gene. *PSMF1* was proposed as a key regulator of human spermatogenesis (Cui *et al.* 2008).

**Table 1.**
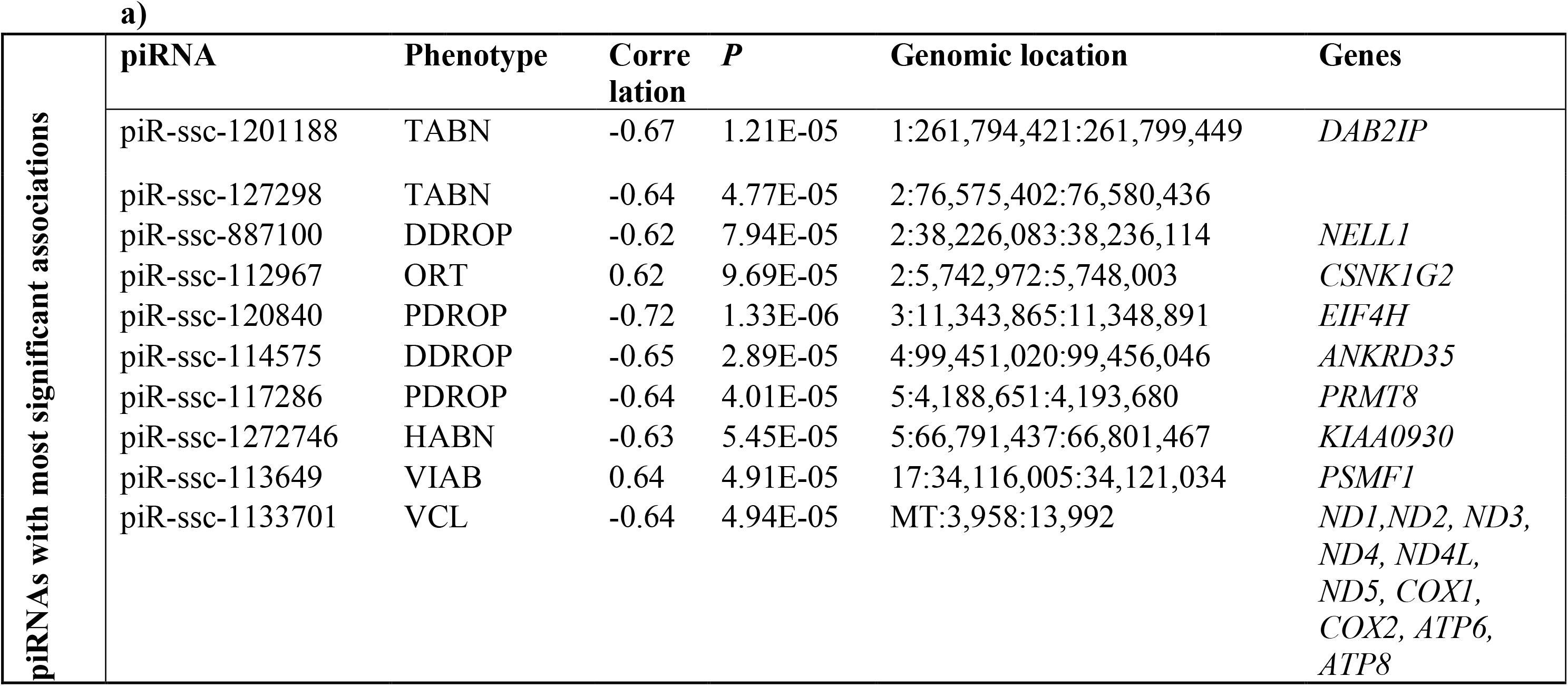
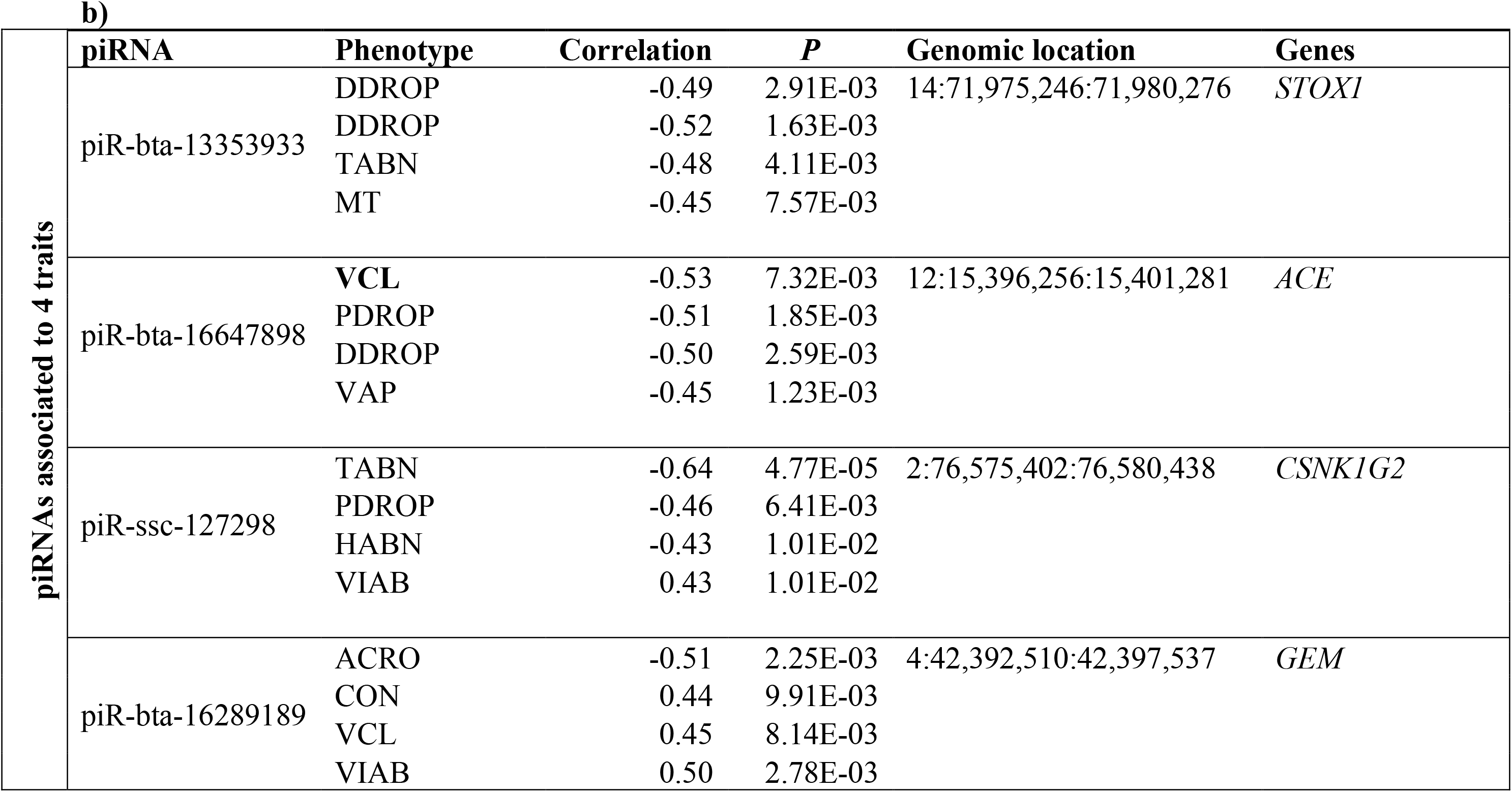
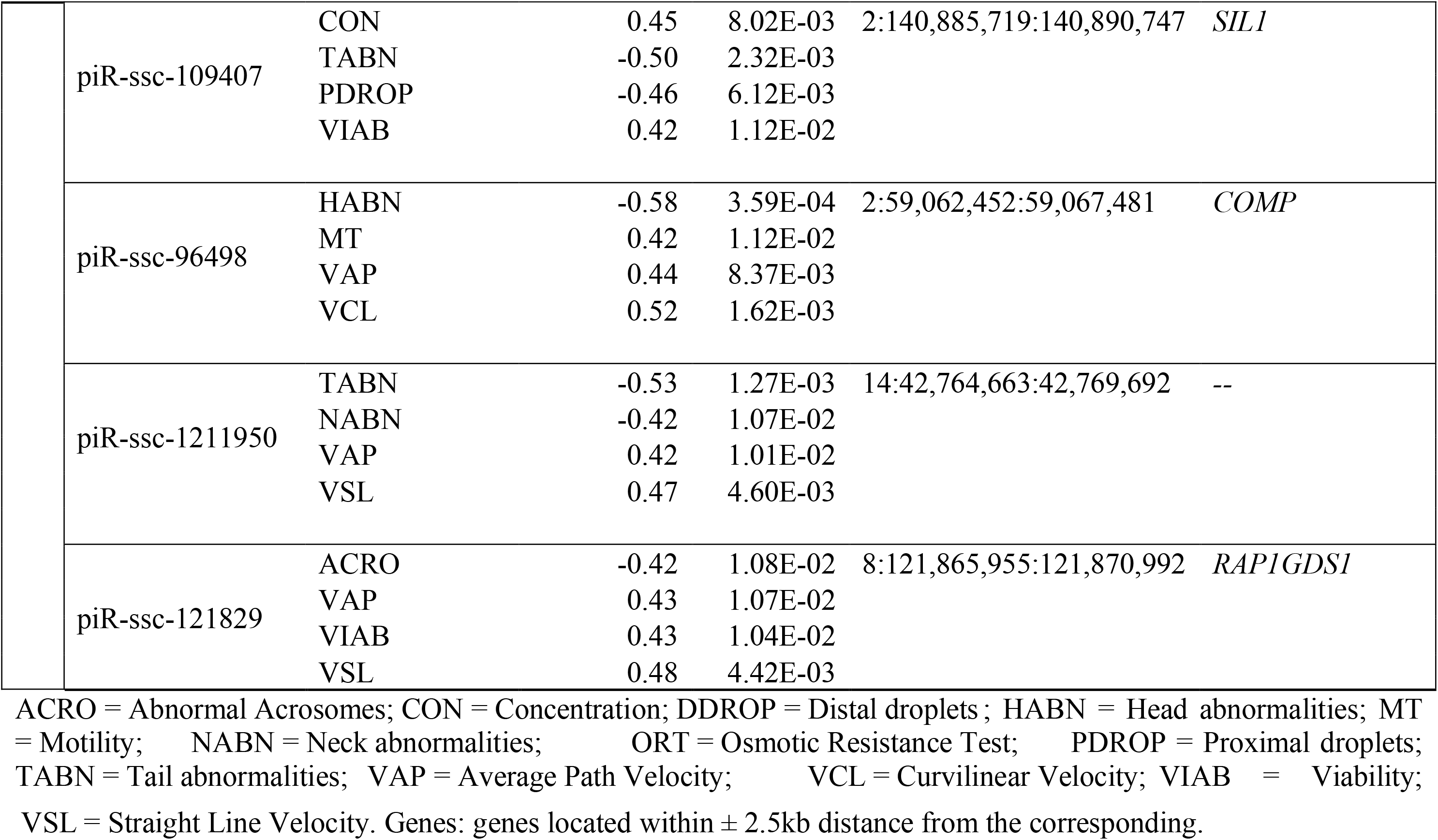
List of piRNAs with most significant (a) or largest number of (b) correlations with semen quality parameters.

The majority of these piRNAs were significantly correlated to only one phenotype (75.7%). On the other side, 8 piRNAs correlated with 4 phenotypes each (Table 1b). Among these, piR-bta-16647898 and piR-ssc-109407 were of great interest. piR-bta-16647898 is negatively correlated with average curvilinear velocity (VCL), average path velocity (VAP), percentage of proximal (PDROP) and distal droplets (DDROP) and it is located near the *ACE* gene (Table 1 b). *ACE* regulates sperm development and functions in human, mice and bull (Fuchs *et al.* 2005; Fujihara *et al.* 2013; Ojaghi *et al.* 2018). piR-ssc-109407, which is positively correlated with VIAB and concentration (CON) and negatively correlated with TABN and PDROP, is located within the *SIL1* gene. Interestingly, *SIL1* seminal plasma protein has been negatively correlated with fertility in bulls (Viana *et al.* 2018).

In conclusion, we characterised the piRNA composition of the porcine sperm and identified several piRNAs that could impact on semen quality traits. These results indicate that piRNAs may hold potential as markers of sperm quality in swine.

## Availability of data

The datasets generated and analysed are available at NCBI’s BioProject PRJNA520978.

## Supporting information

**Table S1.** Description, mean value and Standard Deviation (SD) for each of the 13 semen quality parameters in the 34 samples used in the study.

**Table S2.** Number of human, mouse and swine piRNAs from the piRBase database retrieved, mapped, present in the boar sperm and associated to semen quality.

**Table S3.** Genomic coordinates of the phenotype-associated piRNAs, species of origin and correlated phenotypes. Chr = chromosome; Corr = correlation

**Table S4.** Distance between the phenotype-associated piRNAs and Repetitive Element (RE) classes and number of piRNAs in the vicinity of each RE class. SINE = Short interspersed nuclear element; LINE = Long interspersed nuclear element; DNA/hAT = hAT DNA transposon; LTR = Long terminal repeats.

**Table S5.** Correlation between total piRNA abundance and the 13 phenotypes. Acronym phenotypes can be found in Table S1.

**Table S6.** Biological processes (gene ontology GO term) significantly overrepresented within the set of genes located in the ± 2.5kb window centred at the phenotype associated piRNAs.

**Figure S1.** Relative nucleotide composition of the 6,729 piRNAs with CPM > 1 in more than 75% of the samples from position 1 to position 10.

**Figure S2.** Number of significantly associated piRNAs for each of the 13 phenotypes.

**Figure S3.** Violin plots of piRNA distribution within each Repetitive Element class. SINE = Short interspersed nuclear element; LINE = Long interspersed nuclear element; DNA/hAT = hAT DNA transposon; LTR = Long Terminal Repeat.

## Bibliography

Aravin, A.A., Sachidanandam, R., Bourchis, D., Schaefer, C., Pezic, D., Toth, K.F., Bestor, T., Hannon, G.J. (2008) A piRNA Pathway Primed by Individual Transposons Is Linked to De Novo DNA Methylation in Mice. Molecular Cell 31, 785–799.

Brennecke, J., Aravin, A.A., Stark, A., Dus, M., Kellis, M., Sachidanandam, R., Hannon, G.J. (2007) Discrete Small RNA-Generating Loci as Master Regulators of Transposon Activity in Drosophila. Cell 128, 1089–1103.

Byrne, K., Leahy, T., Mcculloch, R., Colgrave, M.L., Holland, M.K. (2012) Comprehensive mapping of the bull sperm surface proteome. Proteomics 12, 3559–3579.

Cai, X., Yu, S., Mipam, T.D., Yang, F., Zhao, W., Liu, W., Cao, S.Z., Shen, L., Zhao, F., Sun, L., Xu, C., Wu, S. (2017) Comparative analysis of testis transcriptomes associated with male infertility in cattle yak. Theriogenology 88, 28–42.

Cui L, Fang L, Shi B, Qiu S, Ye Y. (2018) Spermatozoa Expression of piR-31704, piR-39888, and piR-40349 and Their Correlation to Sperm Concentration and Fertilization Rate After ICSI. Reproductive Sciences 25, 733–739.

Capra, E., Turri, F., Lazzari, B., Cremonesi, P., Gliozzi, T.M., Fojadelli, I., Stella, A., Pizzi, F. (2017) Small RNA sequencing of cryopreserved semen from single bull revealed altered miRNAs and piRNAs expression between High- and Low-motile sperm populations. BMC Genomics 18, 1–12.

Cui, Y., Zhu, H., Zhu, Y., Guo, X., Huo, R., Wang, X., Tong, J., Qian, L., Zhou, Z., Jia, Y., Lue, Y.H., Hikim, A.S., Wang, C., Swerdloff, R.S., Sha, J. (2008) Proteomic analysis of testis biopsies in men treated with injectable testosterone undecanoate alone or in combination with oral levonorgestrel as potential male contraceptive. Journal of Proteome Research 7, 3984–3993.

Ernst, C., Odom, D.T., Kutter, C. (2017) The emergence of piRNAs against transposon invasion to preserve mammalian genome integrity. Nature Communications 8, 1–9.

Fu, Q., Wang, P.J. (2014) Mammalian piRNAs. Spermatogenesis 4, e27889.

Fuchs, S., Frenzel, K., Hubert, C., Lyng, R., Muller, L., Michaud, A., Xiao, H.D., Adams, J.W., Capecchi, M.R., Corvol, P., Shur, B.D., Bernstein, K.E. (2005) Male fertility is dependent on dipeptidase activity of testis ACE. Nature Medicine 11, 1140–1142.

Fujihara, Y., Tokuhiro, K., Muro, Y., Kondoh, G., Araki, Y., Ikawa, M., Okabe, M. (2013) Expression of TEX101, regulated by ACE, is essential for the production of fertile mouse spermatozoa. Proceedings of the National Academy of Sciences of the United States of America 110, 8111–8116.

Ge, S.Q., Lin, S.L., Zhao, Z.H., Sun, Q.Y. (2017) Epigenetic dynamics and interplay during spermatogenesis and embryogenesis: Implications for male fertility and offspring health. Oncotarget 8, 53804–53818.

Gebert, D., Ketting, R.F., Zischler, H., Rosenkranz, D. (2015) piRNAs from pig testis provide evidence for a conserved role of the Piwi pathway in post-transcriptional gene regulation in mammals. PLoS ONE 10, 1–22.

Di Giacomo, M., Comazzetto, S., Saini, H., De Fazio, S., Carrieri, C., Morgan, M., Vasiliauskaite, L., Benes, V., Enright, A.J., OCarroll, D. (2013) Multiple Epigenetic Mechanisms and the piRNA Pathway Enforce LINE1 Silencing during Adult Spermatogenesis. Molecular Cell 50, 601–608.

Gòdia, M., Estill, M., Castelló, A., Balasch, S., Rodríguez-Gil, J.E., Krawetz, S.A., Sánchez, A., Clop, A. (2019) A RNA-seq analysis to describe the boar sperm transcriptome and its seasonal changes. Frontiers in Genetics 10, 1–14.

Gòdia, M., Swanson, G., Krawetz, S.A. (2018) a A history of why fathers RNA matters. Biology of Reproduction 99, 147–159.

Gòdia, M., Mayer, F.Q., Nafissi, J., Castelló, A., Rodríguez-Gil, J.E., Sánchez, A., Clop, A. (2018) b A technical assessment of the porcine ejaculated spermatozoa for a sperm-specific RNA-seq analysis. Systems Biology in Reproductive Medicine 64, 291–303.

Goh, W.S.S., Falciatori, I., Tam, O.H., Burgess, R., Meikar, O., Kotaja, N., Hammell, M., Hannon, G.J. (2015) PiRNA-directed cleavage of meiotic transcripts regulates spermatogenesis. Genes and Development 29, 1032–1044.

Gunes, S., Kulac, T. (2014) The role of epigenetics in spermatogenesis. Türk Üroloji Dergisi/Turkish Journal of Urology 39, 181–187.

He, Z., Kokkinaki, M., Pant, D., Gallicano, G.I., Dym, M. (2009) Small RNA molecules in the regulation of spermatogenesis. Reproduction 137, 901–911.

Langmead, B. (2010) Aligning Short Sequencing Reads with Bowtie. Current Protocols in Bioinformatics 32, 1–24.

Li, B., He, X., Zhao, Y., Bai, D., Bou, G., Zhang, X., Su, S., Dao, L., Liu, R., Wang, Y., Manglai, D. (2019) Identification of piRNAs and piRNA clusters in the testes of the Mongolian horse. Scientific Reports 9, 1–9.

Liao, Y., Smyth, G.K., Shi, W. (2014) FeatureCounts: An efficient general purpose program for assigning sequence reads to genomic features. Bioinformatics 30, 923–930.

Liu, G., Lei, B., Li, Y., Tong, K., Ding, Y., Luo, L., Xia, X., Jiang, S., Deng, C., Xiong, Y., Li, F. (2012) Discovery of potential pirnas from next generation sequences of the sexually mature porcine testes. PLoS ONE 7, 1–9.

Martin, M. (2011) Cutadapt removes adapter sequences from high-throughput sequencing reads. EMBnet.journal 17, 10.

ODonnell, K.A., Boeke, J.D. (2007) Mighty Piwis Defend the Germline against Genome Intruders. Cell 129, 37–44.

Ojaghi, M., Kastelic, J., Thundathil, J.C. (2018) Testis-specific isoform of angiotensin-converting enzyme (tACE) as a candidate marker for bull fertility. Reproduction, Fertility and Development 30, 1584.

Rosenkranz, D., Han, C.T., Roovers, E.F., Zischler, H., Ketting, R.F. (2015) Piwi proteins and piRNAs in mammalian oocytes and early embryos: From sample to sequence. Genomics Data 5, 309–313.

Salas-Huetos, A., Blanco, J., Vidal, F., Godo, A., Grossmann, M., Pons, M.C., F-Fernández, S., Garrido, N., Anton, E. (2015) Spermatozoa from patients with seminal alterations exhibit a differential micro-ribonucleic acid profile. Fertility and Sterility 104, 591–601.

Savadi-Shiraz, E., Edalatkhah, H., Talebi, S., Heidari-Vala, H., Zandemami, M., Pahlavan, S., Modarressi, M.H., Akhondi, M.M., Paradowska-Dogan, A., Sadeghi, M.R. (2015) Quantification of sperm specific mRNA transcripts (PRM1, PRM2, and TNP2) in teratozoospermia and normozoospermia: New correlations between mRNA content and morphology of sperm. Molecular Reproduction and Development 82, 26–35.

Sun, X. hong, Zhu, Y. ying, Wang, L., Liu, H. ling, Ling, Y., Li, Z. li, Sun, L. bo (2017) The Catsper channel and its roles in male fertility: A systematic review. Reproductive Biology and Endocrinology 15, 1–12.

Tokuhiro, K., Isotani, A., Yokota, S., Yano, Y., Oshio, S., Hirose, M., Wada, M., Fujita, K., Ogawa, Y., Okabe, M., Nishimune, Y., Tanaka, H. (2009) OAZ-t/OAZ3 is essential for rigid connection of sperm tails to heads in mouse. PLoS Genetics 5, 1–7.

Viana, A.G.A., Martins, A.M.A., Pontes, A.H., Fontes, W., Castro, M.S., Ricart, C.A.O., Sousa, M. V., Kaya, A., Topper, E., Memili, E., Moura, A.A. (2018) Proteomic landscape of seminal plasma associated with dairy bull fertility. Scientific Reports 8, 1–13.

Waclawska, A., Kurpisz, M. (2012) Key functional genes of spermatogenesis identified by microarray analysis. Systems Biology in Reproductive Medicine 58, 229–235.

Wang, J., Zhang, P., Lu, Y., Li, Y., Zheng, Y., Kan, Y., Chen, R., He, S. (2019) PiRBase: A comprehensive database of piRNA sequences. Nucleic Acids Research 47, 175–180.

Yan, W. (2017) Male infertility caused by dominant point mutations in the D-box domain reveals a novel role of murine and human PIWI proteins. Biology of Reproduction 96, 1121–1123.

Yang, K., Meinhardt, A., Zhang, B., Grzmil, P., Adham, I.M., Hoyer-Fender, S. (2012) The Small Heat Shock Protein ODF1/HSPB10 Is Essential for Tight Linkage of Sperm Head to Tail and Male Fertility in Mice. Molecular and Cellular Biology 32, 216–225.

Yang, Q., Li, R., Lyu, Q., Hou, L., Liu, Z., Sun, Q., Liu, M., Shi, H., Xu, B., Yin, M., Yan, Z., Huang, Y., Liu, M., Li, Y., Wu, L. (2019) Single-cell CAS-seq reveals a class of short PIWI-interacting RNAs in human oocytes. Nature Communications 10, 1–15.

Zak, L.J., Gaustad, A.H., Bolarin, A., Broekhuijse, M.L.W.J., Walling, G.A., Knol, E.F. (2017) Genetic control of complex traits, with a focus on reproduction in pigs. Molecular Reproduction and Development 84, 1004–1011.

Zhang, Z., Wang, G.L., Li, H.X., Li, L., Cui, Q.W., Wei, C. Bin, Zhou, F. (2012) Regulation of fertilization in male rats by CatSper2 knockdown. Asian Journal of Andrology 14 301–309.

